# Efficient mechano-enzymatic hydrolysis of polylactic acid under moist-solid conditions

**DOI:** 10.1101/2022.11.14.516414

**Authors:** Mario Pérez-Venegas, Tomislav Friščić, Karine Auclair

## Abstract

Quantitative mechano-enzymatic depolymerisation of polylactic acid to lactic acid was achieved at 55°C using the *Humicola insolens* cutinase enzyme in moist-solid reaction mixtures. The resulting lactic acid was easily recovered, and the crude product was pure enough to be used in further synthesis of a value-added compound, a known benzimidazole-based drug precursor. The presented mechano-enzymatic depolymerisation strategy enables the closed-loop recycling of untreated polylactic acid under mild conditions, using a renewable, non-toxic catalyst and producing minimum solvent waste.

## Introduction

The use of biodegradable plastics is an important strategy to counter plastic pollution, a problem fuelled by the growing global production of plastics, reported to be almost 400 million tons in 2020.^1,2^ Polylactic acid (PLA) is one of the best-suited polymers for packaging, textiles, adhesives, and electronic applications.^3^ Besides biocompatibility and thermoplasticity,^3^ PLA is an attractive material due to its higher biodegradability than most commercial fossil- and bio-based plastics. Under environmental conditions, however, the biodegradation of PLA is very slow,^4^ with only minimal weight loss after one year of exposure to marine environments, and a slight degradation after three months of burial in soil (see Supporting Information, **Table S1**).^4^

From a sustainability perspective, biodegradation is a waste destruction option rather than recycling, as it does not take advantage of the energy and the chemistry embedded in the polymer.^5–7^ Recycling is a better alternative, especially when implemented in a closed-loop process, permitting some of the molecular energy and chemistry to be retained in the process. The current thermomechanical approach to recycling of PLA is considered downcycling inasmuch as it causes significant deterioration of the polymer integrity, notably the reduction of its molecular weight and viscosity, while generating a product that cannot be recycled.^8^ Whereas chemical depolymerisation of PLA has been reported, high temperature and pressure conditions are typically needed.^9^ Biocatalytic PLA depolymerisation, predominantly based on the use of hydrolase enzymes, has also been reported.^10^ These methods, however, proceed in aqueous solution and therefore necessitate PLA pre-processing, such as emulsification and composite formation.^10^ Their applicability is generally limited by low efficiency, and/or lack of lactic acid recovery methods.^10^ The interesting properties of PLA (such as biocompatibility, high strength, stiffness, and thermosplasticity), however, warrant further exploration of means to recycle this valuable material (**Table S2**).^1,11,12^

Recently, the use of biocatalysts coupled with mechanical agitation has led to a new development in mechanochemistry, termed mechano-enzymology,^13,14^ which rapidly found applications in areas such as asymmetric synthesis,^15–19^ peptide bond formation,^20,21^ drug synthesis,^18,19,22^ polyester polymerisation,^23^ biopolymer hydrolysis, ^13,24–29^ and even polyethylene terephthalate (PET) plastic depolymerisation.^30–32^ Importantly, early reports in mechano-enzymology established that enzymatic activity can be maintained throughout a continuous milling process,^15–23^ enabling the recovery of the biocatalyst with partial enzymatic activity.^13,15,16,18^ Interestingly, solvent-free, moist-solid state methodologies that incorporate sequential cycles of milling followed by static incubation at a fix temperature, rather than non-stop milling, have also been explored. This novel approach, called reactive aging or RAging,^13^ was initially developed to enhance enzymatic activity towards cellulose, and has been successfully applied and adapted to biocatalytic hydrolysis of other biopolymers such as hemicellulose, lignocellulosic materials, and chitin.^13,24–29^ This work, and the recent report of high crystallinity PET depolymerisation by the cutinase of *Humicola insolens* (HiC) with RAging of moist-solid mixtures,^30–34^ led us now to explore similar conditions for the depolymerisation of PLA. Here, we demonstrate that simple milling followed by aging or RAging reaction conditions at mild temperature enable the clean, quantitative production of lactic acid directly from untreated PLA, using the renewable and non-toxic enzyme HiC. The herein presented mechano-enzymatic method is more efficient than previously described biocatalytic processes, while requiring little aqueous or organic solvent.

## Results and discussion

As mentioned before, the RAging strategy has proven efficient in the depolymerisation of a wide range of biopolymers and of PET plastics. However, considering the simplicity of milling once followed by aging – a processes referred here as MAging, we initially focused on this latter approach.

Our preliminary evaluation of the reactivity of racemic PLA powder (24% crystallinity, typical of materials commonly found in industrial applications), targeted a reaction mixture moistened with a 400 mM Tris-HCl buffer at pH 8. The amount of liquid buffer added to solid PLA was selected to correspond to a liquid-to-solid ratio (*η*) of 1.5 μL/mg, or 6 equivalents of water per monomer unit.^35^ The mixture was placed in a 15 mL poly(tetrafluorethylene) (PTFE, Teflon) milling jar containing one zirconia ball (10 mm diameter, 3.5 g). After addition of the HiC enzyme (Novozyme 51032; 0.65 % w/w relative to PLA), the mixture was mechanically mixed for 5 minutes using an FTS-1000 ball mill operating at a frequency of 30 Hz, followed by static incubation at 55°C for up to 5 days. Under these conditions, HiC was found to catalyse the clean hydrolysis of PLA (a racemic polymer) to racemic lactic acid (2.2±0.1 mg of lactic acid per mL, corresponding to 3.0±0.2% yield). The reaction progress was found to reach a plateau after ca. 2 days of incubation (**Figures 1**, **S1**).

**Figure 1.**
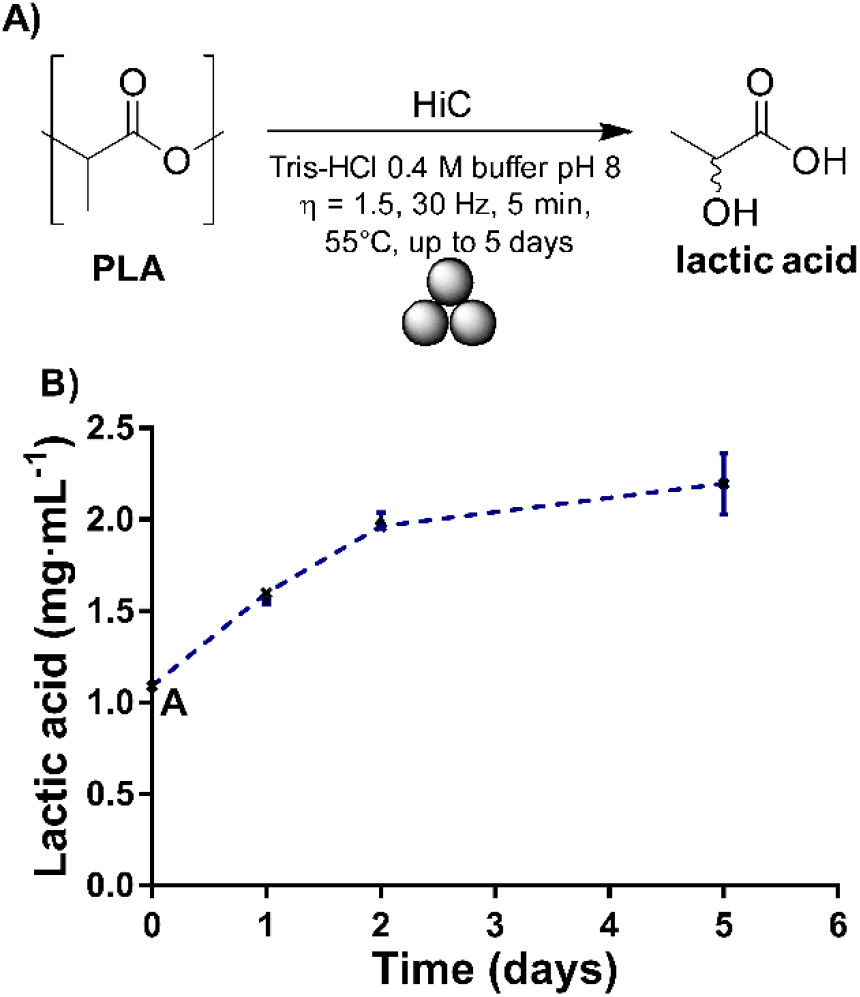
Nature of the reaction studied and initial kinetic data. A) The reaction studied here, showing the initial conditions tested. B) Formation of lactic acid over time under the conditions shown in A), with point A in the graph representing the concentration of lactic acid measured after the milling step only. Experiments were conducted in triplicate; error bars show the standard deviation of the mean.

Following these encouraging results, optimization of the process was undertaken by sequentially varying a range of parameters known to affect mechano-enzymatic reactions (*e.g*., solid-to-liquid ratio *η*, buffer pH, aging temperature, enzyme loading, and effect of additives).^36^ Increasing *η* from 1.5 to 4.5 μL/mg (corresponding to 18 equivalents of water per monomer lactic acid), was found to enhance the yield of lactic acid to 10.9±0.4% (**Figure 2A**). The use of higher *η*-values did not lead to further beneficial effect on the reaction yield.

**Figure 2.**
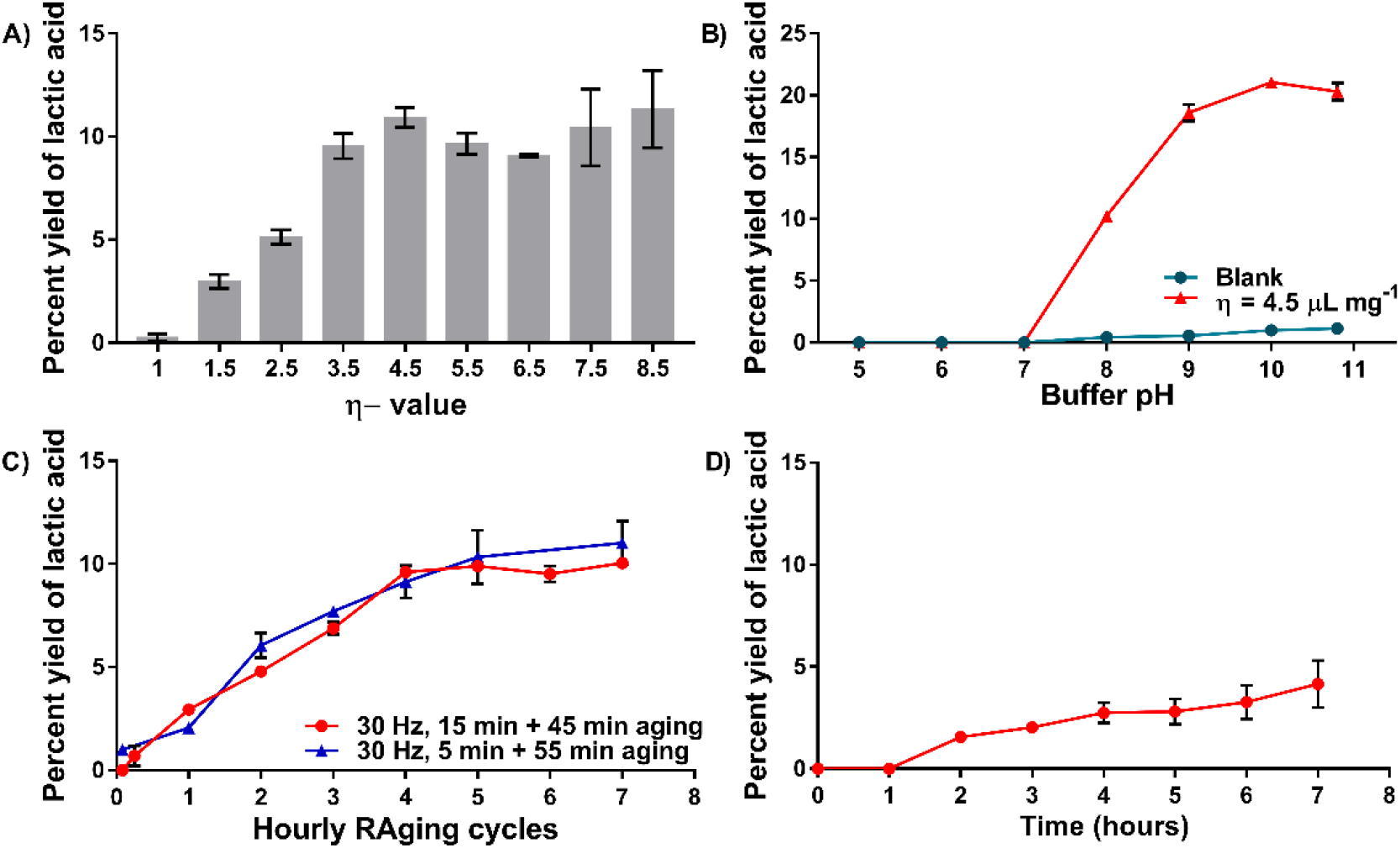
Optimization of HiC-catalyzed (0.65% w/w loading) PLA depolymerisation using the conditions given in Figure 1 after 3 days of reaction, unless noted otherwise. A) Effect of the liquid-to-solid ratio (*η*) of the reaction mixture on the yield of lactic acid. B) Effect of varying the buffer pH (*η* = 4.5 μL·mg^−1^) at a buffer concentration of 0.4 M (sodium citrate buffer for pH 5-6, Tris-HCl buffer for pH 7-9, sodium borate buffer for pH 10, sodium carbonate buffer for pH 10.8). Blank reaction mixtures were treated the same way, but were missing the enzyme. C) Reaction progress under RAging conditions over 7 h (15 min of milling + 45 min of aging at 55°C, or 5 min of milling + 55 min of aging at 55°C) in 0.4 M sodium borate buffer at pH 10 and *η* = 4.5 μL·mg^−1^. D) Reaction progress under milling + aging conditions (MAging) over 7 h (15 min of milling at 30 Hz in 0.4 M sodium borate buffer at pH 10 and *η* = 4.5 μL·mg^−1^, incubation at 55°C). Experiments were conducted in triplicate; error bars show the standard deviation of the mean.

We worried that the high water solubility of the lactic acid product and its acidity (p*K_a_* of 3.86), may interfere with the enzymatic process by acidifying the mixture. To evaluate the effect of lactic acid on the mechano-enzymatic transformation, lactic acid was added to the mixture at the start of the reaction. We observed that the reaction yield was considerably reduced in the presence of product, with only 3% of fresh lactic acid being produced after 3 days of incubation at 55°C (data not shown). Measurement of the pH of the reaction mixture revealed that the added lactic acid contributed to lowering the pH from 10 to 7.2 – a pH at which HiC has negligible activity (**Figure 2B**). Conversely, a higher yield of lactic acid (21.0±0.4%) was obtained when the initial pH of the buffer was increased from 8 to 10 by using a 0.4 M sodium borate buffer to *η* = 4.5 μL/mg (**Figure 2B**).

Next, we focused on the temperature of the static incubation process. Decreasing the incubation temperature from 55°C to 25°C or 45°C reduced the yield of lactic acid from 21.0±0.4% to 7±3% and 19±1%, respectively (**Figure S2**). Moreover, increasing the temperature of the static incubation step to 65°C resulted in a negligible increase in the yield of lactic acid to 21.2±0.3% (**Figure S2**). Similarly, decreasing the enzyme loading to 0.21% and 0.43% by weight led to a slight decrease in the lactic acid yield, to 17.1±0.6% and 18.8±0.3%, respectively. To our surprise, increasing the enzyme loading to 1.3% w/w was also found to decrease the lactic acid yield to 17.2±1.4% (**Figure S2**). We also investigated the possibility that small volumes of an organic solvent, such as methanol (MeOH), ethanol (EtOH), iso-propyl alcohol (*i*-PrOH), acetonitrile (CH_3_CN), dichloromethane (CH_2_Cl_2_) or toluene might lead to reaction acceleration, but this did not prove successful (**Figure S2**). Accordingly, optimization of the reaction was further pursued with 0.65% w/w HiC at an aging temperature of 55°C without including an organic solvent additive.

Next, we explored the conversion of PLA to lactic acid by RAging, *i.e*., through a process in which the moist-solid reaction mixture is exposed to alternative periods of brief milling and aging.^13,24–31^ Notably, periodic, hourly repeated milling of the reaction mixture for 5 or 15 minutes led to a noticeable improvement in the lactic acid yield (10.0±0.3% yield of lactic acid in 4 hours, **Figure 2C**), with respect to the MAging process (2.7±0.5% of lactic acid after 4 hours of aging, **Figure 2D**). Conversion of PLA to lactic acid, however, did not increase significantly with further RAging cycles, possibly due to accelerated enzyme denaturation induced by repeated miling processes.^30^ Moreover, repeating the milling process on a daily basis, using identical overall conditions (0.65% w/w HiC, 0.4 M sodium borate buffer at pH 10, *η* = 4.5 μL/mg, static incubation at 55°C) was less promising (**Figure S3**). Consequently, to minimize enzyme denaturation, most subsequent experiments used milling only at the beginning of the process.

To further overcome acidification of the reaction medium by the lactic acid product, we next increased the buffer concentration (**Figure 3**). As anticipated, this was greatly beneficial to the process, with the reaction yield rising to 44.8±0.8% at a buffer molarity of 1 M (pH 10, *η* = 4.5 μL/mg) in a process consisting of milling for 5 minutes, followed by three days of static incubation at 55°C (**Figure 3**). Nevertheless, after 3 days of reaction, formation of the lactic acid product had again acidified the reaction mixture to pH 7.3. Importantly, control experiments with only PLA and the buffer (1 M, pH 10) showed negligible hydrolysis, implying that the 44.8% yield is due to HiC activity, rather than basic hydrolysis.

**Figure 3.**
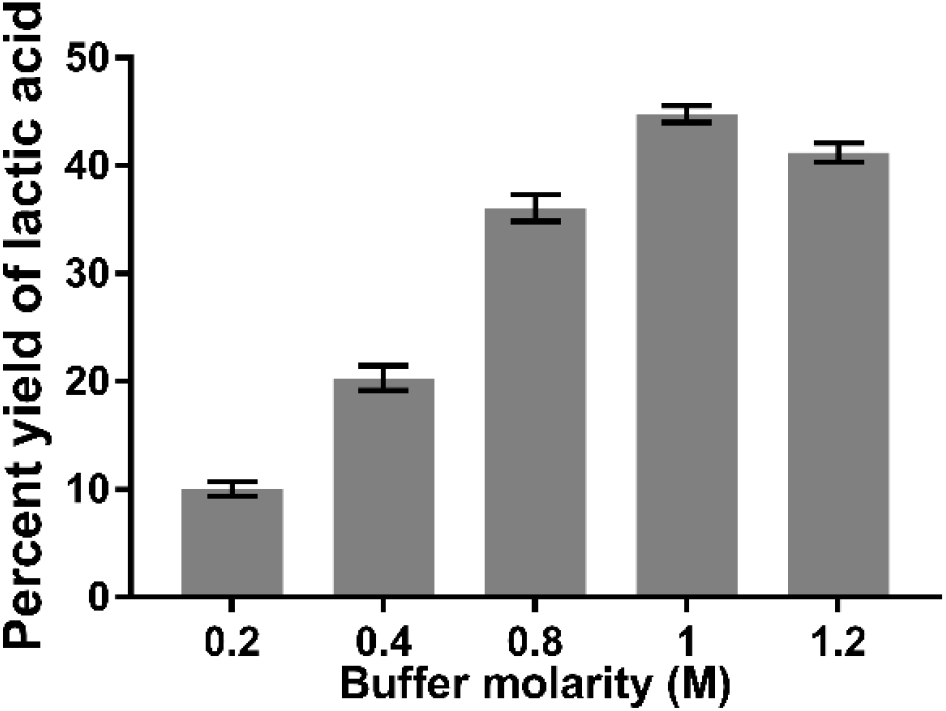
Effect of buffer molarity (sodium borate pH 10, *η* = 4.5 μL·mg^−1^) on the lactic acid yield from PLA using HiC (0.65% w/w) with mixtures treated to 15 minutes of milling (30 Hz) followed by 3 days of static incubation at 55°C. Experiments were conducted in triplicate; error bars show the standard deviation of the mean.

Buffers of even higher molarity were also tested as liquid additives for the mechanically-activated moist-solid reaction mixtures. Due to the poor solubility of sodium borate in water, it was necessary to switch to Tris and glycine buffers to reach 2-3 M (**Figure 4**). In 2 M glycine buffer at pH 10, a 35±1% yield of lactic acid was obtained after 3 days (39±2% yield after 7 days). Notably, a 3 M Tris buffer at pH 9, afforded 65±3% of lactic acid in 4 days. Such a high yield and selectivity for lactic acid is unprecedented without pre-processing of the PLA substrate (**Table S2**).^1,9,12^ Moreover, this mechano-enzymatic depolymerisation proved to be as efficient as a similar process in slurry conditions under continuous stirring at 300 rpm (**Figure S4**). Milling the mixture hourly also gave similar results (**Figure S5**). Importantly, control experiments in the absence of enzyme allowed buffer-catalysed hydrolysis to be ruled out.

**Figure 4.**
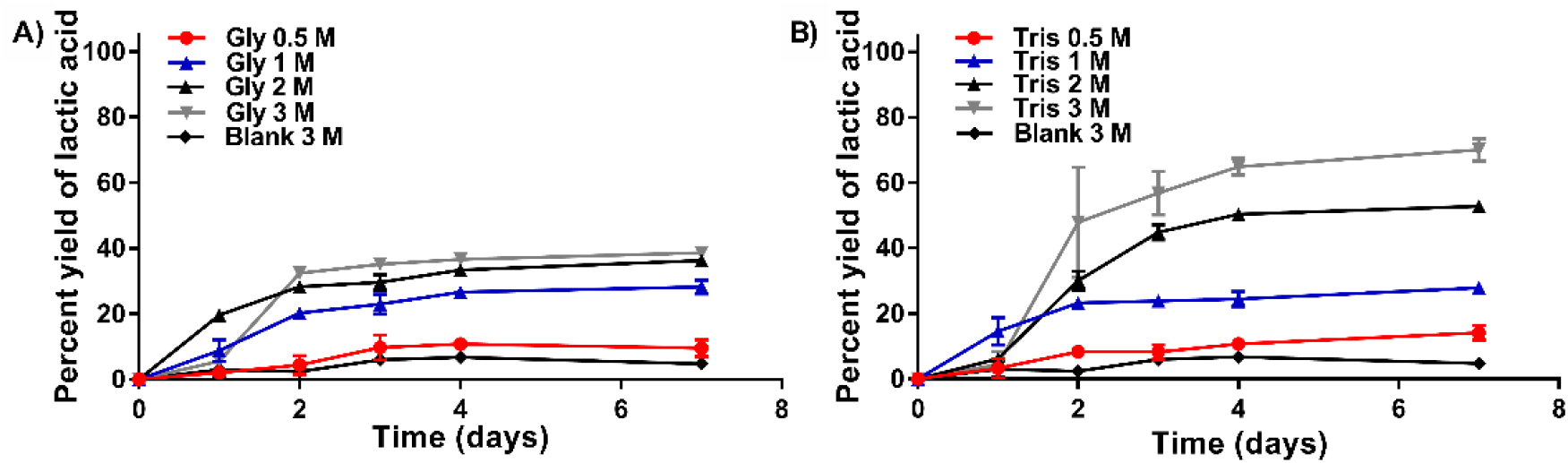
Effect of further increasing the buffer molarity (*η* = 4.5 μL·mg^−1^) on the production of lactic acid from PLA, in a reaction catalysed by HiC (0.65% w/w) with milling for 15 minutes at 30 Hz followed by static incubation at 55°C for variable periods. The buffers used were A) 0.5-3 M glycine buffer at pH 10, and B) 0.5-3 M Tris buffer at pH 9. The blank reactions mixtures contained no enzyme. Experiments were conducted in triplicate; error bars show the standard deviation of the mean.

Unexpectedly, a small extent of PLA aminolysis induced by HiC was also observed in high molarity buffers (*η* = 4.5 μL·mg^−1^, incubated at 55°C for up to 4 days, **Figures S6, S7, S11-S15**), providing amide products from the coupling of lactic acid and glycine (*N*-lactylglycine, *rac*-**2**, **Figure 5A**), or of lactic acid and Tris (*N*-[Tris(hydroxymethyl)methyl]lactamide, *rac*-**4**, **Figure 5A**). The amide products were quantified by HPLC, revealing a total depolymerisation extent of 42±1% in 2 M glycine buffer, and of 70% in 3 M Tris buffer (**Figure 5**). It was also established that lactic acid is not an intermediate in PLA aminolysis since introducing fresh lactic acid to the reaction mixture produced no visible increase in amide product (**Figure S8**).

**Figure 5.**
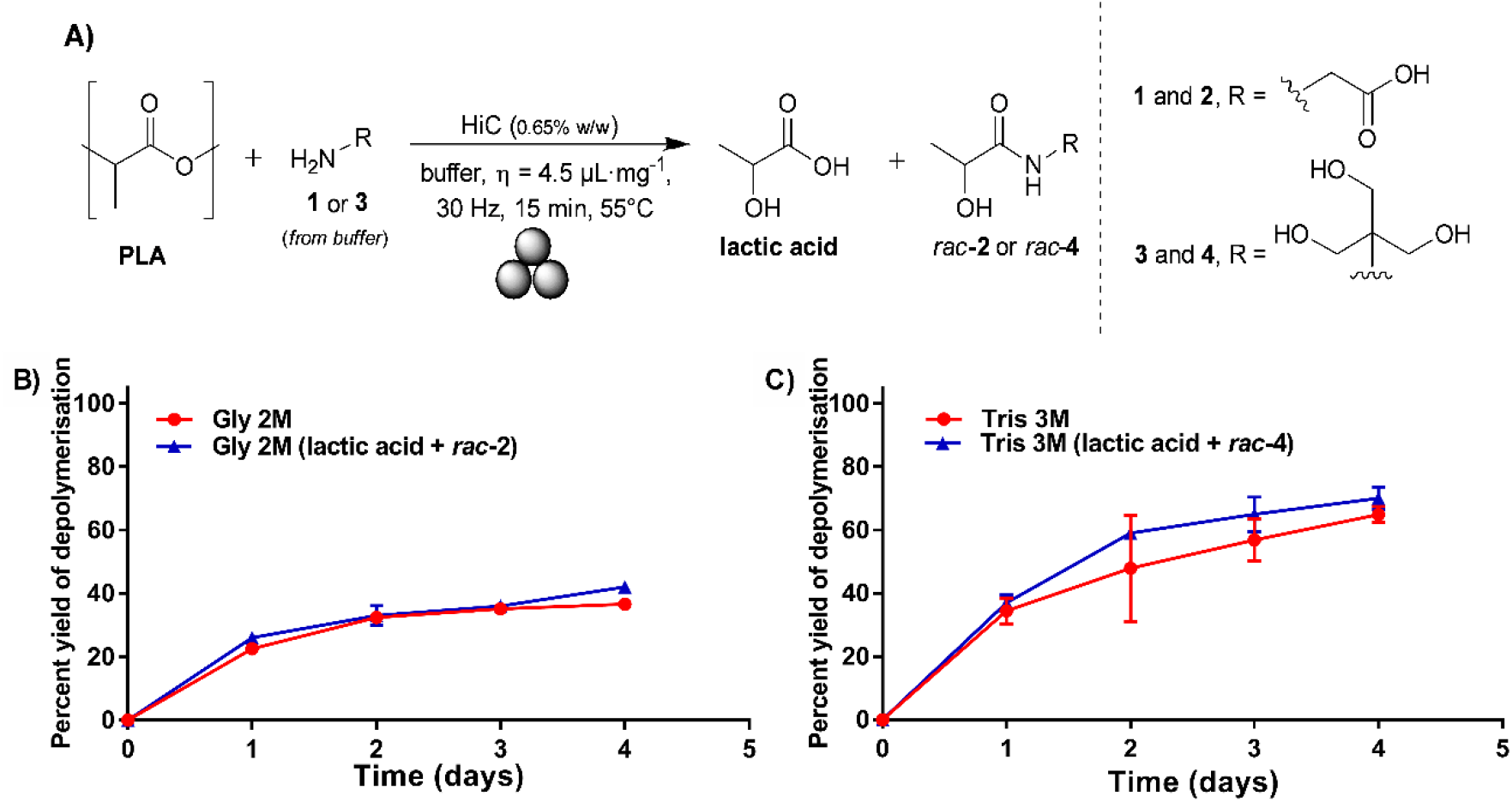
PLA depolymerisation (red: lactic acid alone, or blue: lactic acid + amide) from HiC-catalyzed (0.65% w/w) reactions in moist-solid conditions (*η* = 4.5 μL·mg^−1^) treated to milling for 15 minutes at 30 Hz followed by static incubation at 55°C for variable periods. The buffers used were B) 2 M glycine buffer at pH 10 or C) 3 M Tris buffer at pH 9. Experiments were conducted in triplicate; error bars show the standard deviation of the mean.

Based on the control experiments, we can identify HiC (specifically, Novozyme 51032) as the only promoter of the formation of the amide product. Amide formation catalyzed by cutinases has only been reported for *Fusarium solani pisi* cutinase and an engineered *Humicola insolens* cutinase;^37–40^ the amidase/protease activity of commercial Novozyme 51032 (*Humicola insolens* cutinase expressed in *Aspergillus oryzae*) has not been reported. To get insight into the protease activity of Novozyme 51032, assays using an *o*-nitrophenyl amide model substrate (**Scheme S1**) were performed in 2 M glycine or 3 M Tris buffer at different pH values (**Figure S9**). Indeed, Novozyme 51032 was found to show hydrolytic activity against the tested amide, consistent with the observed HiC-catalyzed aminolysis of PLA with buffer amines.

To further optimize the depolymerisation process, it was envisaged that adding a second batch of buffer just before the reaction plateau is reached (**Figure 4**, 2 days of incubation at 55°C) could further help neutralize the lactic acid product and be beneficial to the process. This was successful, as shown by the quantitative depolymerisation of PLA observed when one-third of the initial amount of 3 M Tris buffer was replenished after two days of incubation at 55°C (**Figure 6B**), simultaneously increasing the *η*-value to 6 μL/mg, with minimal aminolysis of lactic acid (6% yield of **4**, **Figure 6B**). A significant, but less pronounced improvement of the lactic acid yield to 58% was also obtained when the same strategy was applied to reactions using the 2 M glycine buffer (**Figure 6A**).

**Figure 6.**
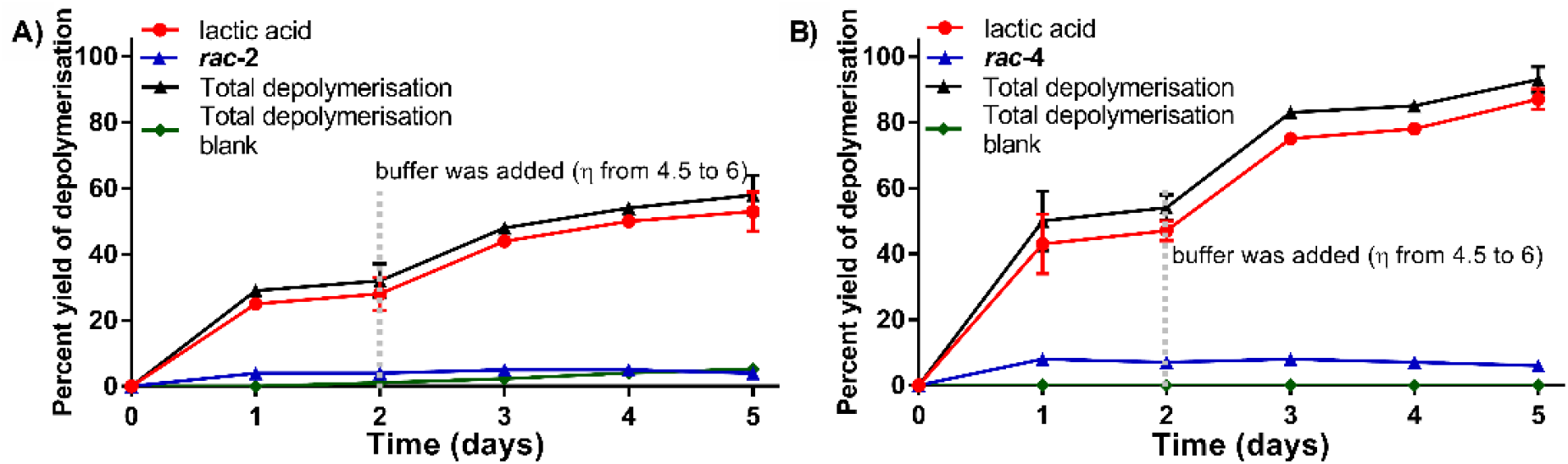
Effect of adding a second batch of buffer (which also translates into increasing the *η*-value from 4.5 to 6 μL/mg) after milling for 15 minutes at 30 Hz and two days of incubation at 55°C for HiC-catalyzed (0.65% w/w) reactions using: a) 2 M glycine at pH 10; b) 3 M Tris pH 9 buffer. Experiments were conducted in triplicate; error bars show the standard deviation of the mean.

The beneficial effect of adding a second buffer batch is likely the result of buffer neutralization. This is further supported by a lack of yield increase when MilliQ® water (to the same *η*-value) is added after two days of incubation instead of the high molarity buffer (**Figure S10**). Additional control experiments under identical conditions but lacking the enzyme showed no detectable hydrolysis. It is envisaged that the addition of a second batch of buffer could be avoided with the use of an acidophilic enzyme variant for example.

Lactic acid finds application in the preparation of foods (as a flavor and acidulant), pharmaceuticals (implants, pills, and sutures), cosmetics (hygiene and aesthetic products), and chemicals (synthetic precursor and building block for the preparation of acids, esters, biosolvents, etc.).^41,42^ With this in mind, we explored the possibility of using the crude lactic acid generated from enzymatic PLA depolymerisation as a substrate in the synthesis of a valuable benzimidazole intermediate, *rac*-**5**, which has been used in the preparation of the anthelmintic drug thiabendazole.^43^ Thus, after quantitative enzymatic depolymerisation of PLA under moist-solid conditions (**Figure 6**, HiC loading of 0.65% w/w, 3 M Tris buffer at pH 9 at *η* = 4.5 μL·mg^−1^, milled for 15 minutes at 30 Hz and incubated at 55°C, with addition of a second batch of buffer after 2 days of incubation, to a final *η* = 6 μL·mg^−1^), the lactic acid product was collected by solid-liquid extraction using a small volume of water (2 mL for 300 mg of substrate), and this aqueous solution was mixed with *o*-phenylenediamine before acidification (HCl 6 N) to pH 1, to form the desired *rac*-**5** in 60% yield (**Scheme 1**), comparable to the same reaction performed with fresh lactic acid (65% yield overall).

**Scheme 1.**
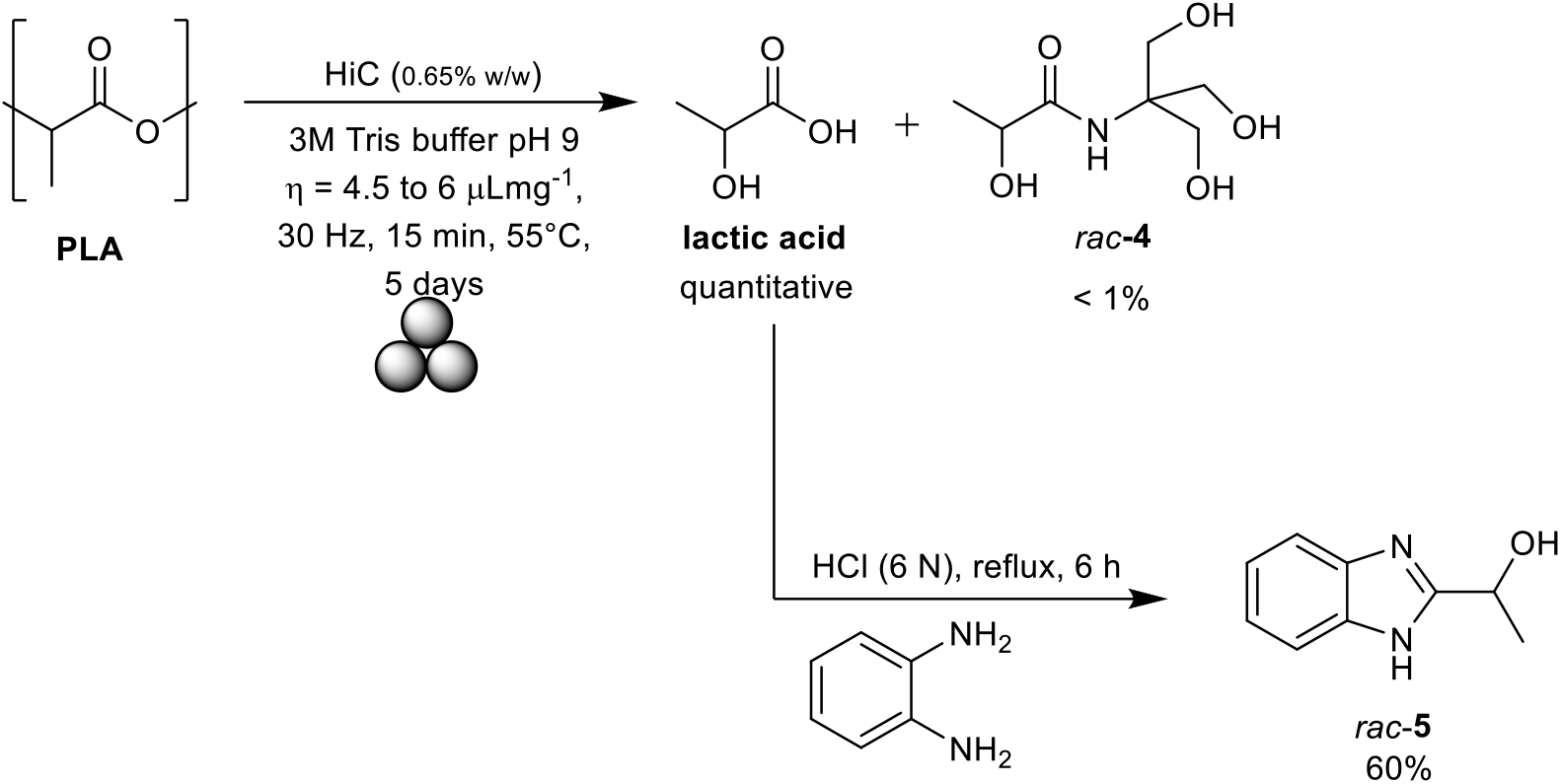
Synthesis of *rac*-**5**, an important intermediate in the synthesis of thiabendazole, from PLA via a chemoenzymatic cascade involving first mechano-enzymatic depolymerisation of PLA, followed by reaction with *o*-phenylenediamine under acidic conditions. The conversion of PLA to lactic acid was quantified by HPLC and the yield of *rac*-**5** by NMR.

## Conclusions

We have demonstrated that mechano-enzymatic depolymerization in moist-solid reaction mixtures is a promising methodology for PLA depolymerization, a process compatible with closed-loop PLA recycling.^14,44,45^ The method established here takes place under mild conditions, with a renewable, non-toxic catalyst and minimal solvent use. The reaction is 200% faster than biocatalytic processes under standard aqueous conditions using untreated PLA.^4,10^ Furthermore, product recovery is simple, and the crude product is sufficiently clean for use as a starting material in the synthesis of a benzimidazole-based drug precursor. Hence, we envisage that this methodology could benefit the revalorization of post-consumer synthetic polymers, such as PLA, that are commonly used in single-use products (*e.g*., cups, bottles, and food containers), devices that have recently been introduced in medicine (including surgical tools, implants, and casts),^46^ and in other applications (*e.g*., PLA filaments for 3D printers, antennas, microwave absorbers).^47,48^

## ASSOCIATED CONTENT

### Supporting Information

General methods, ^1^H and ^13^C NMR of lactic acid, *rac*-**2**, *rac*-**4**, and *rac*-**5**.

