## Supplemental files for "Efficient mechano-enzymatic hydrolysis of polylactic acid under moist-solid conditions"

### Table of contents

| | |
|---|---|
| **Materials** | S2 |
| **Equipment** | S2 |
| **Methods** | S3 |
| *Method to calculate the percentage yield of lactic acid* | S3 |
| *General method for the HiC-catalyzed depolymerisation of PLA under moist-solid conditions* | S3 |
| *Method to verify if lactic acid may inhibit the reaction* | S3 |
| *Measuring the effect of temperature, enzyme loading, and organic solvent on the catalytic activity of HiC in PLA depolymerisation reactions* | S4 |
| *General method for the HiC-catalyzed depolymerisation of PLA under RAging conditions* | S5 |
| *General method for the HiC-catalyzed depolimerisation of PLA under slurry conditions ($\eta$=10)* | S6 |
| *General method for HiC-catalyzed depolimerisation of PLA under RAging conditions using 2M glycine buffer* | S7 |
| *PLA aminolysis promoted by the use of high molarity buffer and catalyzed by HiC under moist-solid conditions* | S8 |
| *Control experiments to rule out lactic acid as promoter in PLA aminolysis under moist-solid conditions* | S9 |
| *Spectrophotometric assay to quantify the protease activity of HiC using N-(4-nitrophenyl) butyramide as substate* | S10 |
| *General method for HiC-catalyzed depolymerisation of PLA with a second addition of buffer after 2 days* | S10 |
| *Chemoenzymatic synthesis of rac-**5** from PLA* | S11 |
| *Characterization of lactic acid, amides products rac-2 and rac-4, and* rac-*5* | S12 |
| **Table S1.** Reported PLA biodegradation experiments mimicking natural conditions | S17 |
| **Table S2.** Reported biocatalytic PLA depolymerisation reactions | S18 |
| **References** | S19 |

### Materials

Unless specified otherwise, DL-lactic acid (ca. 90% purity) was purchased from Sigma-Aldrich and used without further purification. The water used was of MilliQ® quality with a specific resistance of 18.2 MΩ·cm at 25°C. HPLC grade acetonitrile was purchased from Thermo Fisher Scientific (Waltham, MA, US). NMR samples were prepared with $D_2O$ and MeOH-*d*4 purchased from Cambridge Isotope Laboratories, Inc (Tewksbury, MA, US).

The Tris buffer was prepared from Tris-HCl (Chem-Impex; Wood Dale, IL, US), or Tris base (Chem-Impex; Wood Dale, IL, US), the sodium borate buffer was prepared from boric acid (EMD Chemicals; Darmstadt, Germany), and the glycine buffer was prepared from glycine (Chem-Impex; USP 99.6%, Wood Dale, IL, US).

Novozyme®51032, *Humicola insolens* cutinase expressed in *Aspergillus oryzae*, was purchased from Strem Chemicals, Inc. (Newburyport, MA, US; LOT 36025400).

Polylactic acid (PLA) granules were purchased from Goodfellow (Huntingdon, England). The PLA powder was obtained by freezing the PLA granules with liquid nitrogen before ball milling for 5 minutes at 30 Hz using a stainless-steel milling jar (15 mL) and one stainless steel ball (1.5 cm diameter) for at least three times, powder with a particle size below 250 μm was used in the reactions.

### Equipment

A Mettler Toledo AB135-S/FACT DualRange analytical balance (linearity 0.2 mg, readability 0.01 mg/0.1 mg), Mettler Toledo XP105 DeltaRange analytical balance (linearity 0.15 mg, readability 0.01 mg/ 0.1 mg), and VWR pre-calibrated pipettors were used for measuring out the reaction components. Ball milling was performed using a FormTech Scientific FTS1000 shaker mill. Sleeved PTFE jars (15 mL) with one $ZrO_2$ ball (10 mm diameter) were used for milling. Static incubation at 55°C was performed in a Fisher Scientific Oven, whereas static incubation (shaking turned off) at other temperatures used a ShelLab shaking incubators, VWR and Ohaus tabletop shaker incubators, or a Thermolyne Oven Series 900. All reactions were performed in triplicate. Error bars in graphs represent the standard deviation.

A Branson 2510 sonicator was used to prepare samples for HPLC analysis, and Thermo Scientific Sorvall Legend Micro21 and Lynx 6000 centrifuges were used for centrifugation, with volumes of up to 2 mL and to 25 mL, respectively. Lyophilization was achieved using a Labconco FreeZone 1 Liter Benchtop Freezer Dry System.

$^{1}H$ and $^{13}C$ solution-state NMR spectra were acquired on a Varian Mercury 300 MHz spectrometer. Chemical shifts are reported in ppm and referenced to the solvent residual signal; $D_2O$ ($\delta$ = 4.79 ppm) MeOH ($\delta$ = 3.31 ppm). NMR splitting patterns are reported as singlet (s), doublet (d), quartet (q), multiple signal (m). Spectra were analyzed and plotted using MestReNova 14.2.2 software.

HPLC analysis was conducted on an Agilent 1100 series equipped with a quaternary pump, autosampler, and multiwavelenght UV-vis detector. Separation was achieved on a Phenomenex Luna® 5 μm C18(2) 100 Å, 4.6 x 250 mm column with an isocratic 99(A):1(B) mobile phase, where A is 0.1% formic acid in water and B is acetonitrile. The injection volume was set to 10 μL, and the flow rate was 0.7 mL min$^{-1}$ with detection at 210 nm. Commercial DL-lactic acid was used as a standard dissolved in water in concentrations ranging from 2.5 to 10 mg/mL, and eluted at a retention time of 7.4 minutes. Research samples were resuspended in water, sonicated, centrifuged, and filtered through a syringe using a 0.22 μm nylon filter (Chromspec, Brockville, ON, Canada) before HPLC analysis.

### Methods

#### *Method to calculate the percentage yield of lactic acid*

The yield of lactic acid from PLA depolymerisation reactions was calculated using the calibration curve below (**Figure S1**), which is based on HPLC peak area (AUC). The goodness of fit was $R^2$ = 0.9855, given the equation AUC = 471*[lactic acid] + 132.

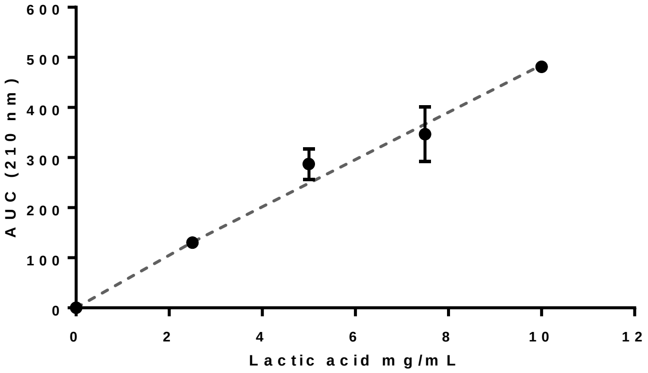


**Figure S1**. Lactic acid calibration curve.

#### *General method for the HiC-catalyzed depolymerisation of PLA under moist-solid conditions*

The PLA powder (300 mg, 4.16 mmol) was placed in a 15 mL sleeved PTFE milling jar containing one $ZrO_2$ ball (10 mm diameter). After addition of the commercial enzyme preparation (300 μL, 1.95 mg protein) and buffer at the desired pH (150 μL for $\eta$ = 1.5 μl mg$^{-1}$, 1050 μL for $\eta$ = 4.5 μl mg$^{-1}$), the reaction mixture was treated to ball milling at 30 Hz for 5 or 15 minutes, before static incubation at 55°C (unless stated otherwise). Samples of the reaction mixture (15-40 mg) were collected for HPLC analysis just after milling, after 1, 3, 5, and 7 days. Experiments were performed in triplicate.

#### *Method to verify if lactic acid may inhibit the reaction*

Whether the presence of lactic acid in the mixture could inhibit the reaction was tested as follows. The PLA powder (125 mg, 1.73 mmol) was placed in a 15 mL sleeved PTFE milling jar containing one $ZrO_2$ ball (10 mm diameter), before addition of the commercial enzyme preparation (125 μL, 0.80 mg protein), 0.4 M sodium borate buffer at pH 10 (437.5 μL, $\eta$ = 4.5 μl mg$^{-1}$), and fresh lactic acid.

The amount of lactic acid added was calculated to match the yield of depolymerisation reached when using 0.4 M sodium borate buffer at pH 10 (21%, **Figure 2B**), which is equivalent to 31.5 mg of lactic acid (or 29 μL). The reaction mixture containing fresh lactic acid was first milled at 30 Hz for 15 minutes, before dividing it into 15-40 mg aliquots and incubating at 55°C for 3 days. The content of lactic acid of each sample was measured by HPLC, and the reaction yield was calculated after subtracting the amount of fresh lactic acid added at the start of the reaction. The reaction mixture was also washed with water to collect the insoluble PLA, which was quantified after lyophilization.

The effect of lactic acid on the pH of the buffer was also measured by adding fresh lactic acid directly into the buffer alone. This was achieved by mixing 5 mL of 0.4 M sodium borate buffer at pH 10 and 331 μL of lactic acid (matching the 437.5 μL to 29 μL ratio used above) and measuring the pH of the solution.

#### ***Measuring the effect of temperature, enzyme loading, and organic solvent on the catalytic activity of HiC in PLA depolymerisation reactions***

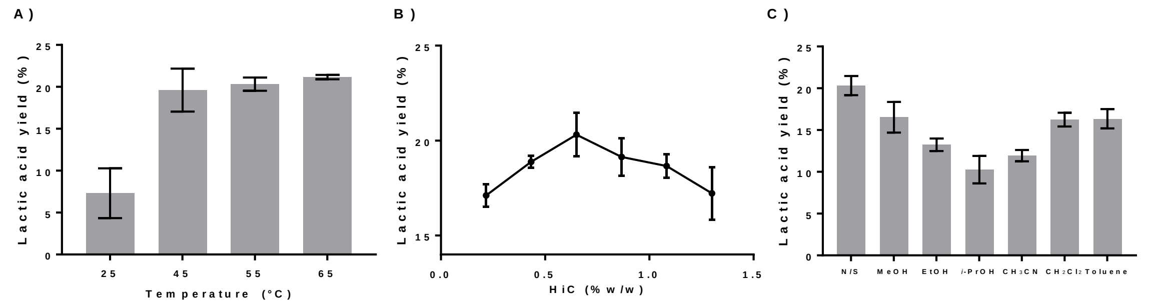


**Figure S2**. Effect of varying the incubation temperature (A) or the enzyme loading (B), and of adding small volumes (150 μL) of organic solvent (C) on the yield of lactic acid from PLA; N/S: No organic solvent was added. The standard conditions used PLA powder (150 mg), 0.4 M sodium borate buffer at pH 10 ($\eta$ = 4.5 μL $mg^{-1}$), and 0.65 %w/w HiC, milled at 30 Hz for 15 minutes and incubated at 55°C for 3 days.

#### *General method for HiC-catalyzed depolymerisation of PLA under RAging conditions*

The PLA powder (300 mg, 4.16 mmol) was placed in a 15 mL unsleeved PTFE milling jar containing one $ZrO_2$ ball (10 mm diameter). After the addition of commercial HiC enzyme (300 μL, 1.95 mg protein) and 0.4 M Tris buffer at pH 10 (1050 μL, $\eta$ = 4.5 μl $mg^{-1}$), the mixture was milled at 30 Hz for 15 minutes either hourly or daily, between periods of static incubation at 55°C. Samples (100-200 mg) were collected regularly for HPLC analysis. Experiments were performed in triplicate. Results are presented on **Figure 2C** of the main manuscript for hourly milling and on **Figure S3** for daily milling.

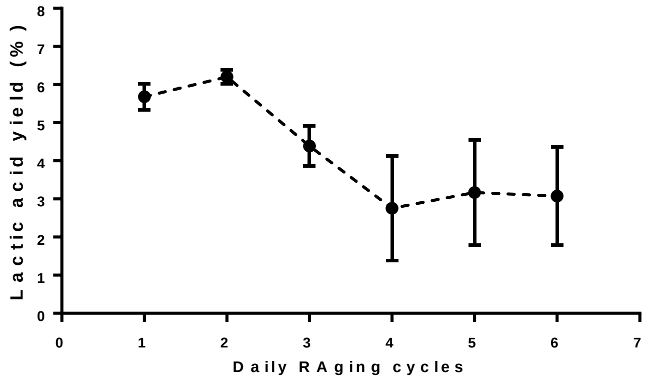


**Figure S3**. Effect of daily milling (30 Hz, 15 minutes) on the yield of lactic acid for the HiC-catalyzed depolymerisation of PLA. The reactions were performed at $\eta$ = 4.5 $\mu L \cdot mg^{-1}$ using 0.4 M sodium borate buffer at pH 10, with 300 mg of PLA, 0.65%w/w HiC, and static incubation at 55°. Experiments were conducted in triplicate; error bars show the standard deviation.

***General method for HiC-catalyzed depolymerisation of PLA under slurry conditions ($\eta$ = 10)***

The PLA powder (100 mg, 1.38 mmol) was placed in a 5 mL round bottom flask. After addition of the commercial enzyme preparation (100 μL, 0.65 mg protein) and 2 M glycine buffer at pH 10 or 3 M Tris buffer at pH 9 using $\eta$ = 10 μl $mg^{-1}$ (900 μL), the reaction mixture was placed on a tabletop orbital shaker and was incubated at 55°C with shaking at 300 rpm. Reaction samples (25 μL) were collected after 1, 2, 5, 7, and 9 days of incubation, and analyzed by HPLC. The experiments were performed in triplicate. The results are shown in **Figure S4**.

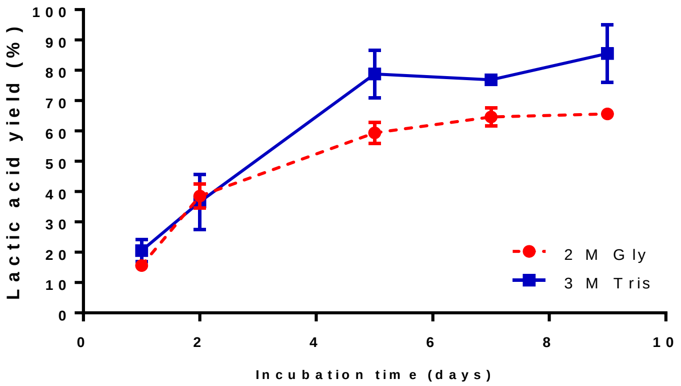


**Figure S4**. HiC-catalyzed depolymerisation of PLA under slurry conditions ($\eta$ = 10 μL·$mg^{-1}$). Reaction mixtures containing PLA (100 mg), HiC (100 μL, 0.65% w/w), and buffer (2 M glycine pH 10 or 3 M Tris pH 9 buffer, 900 μL) were incubated in a tabletop orbital shaker at 55°C and 300 rpm. Reaction samples (25 μL) were extracted at the indicated time and were analyzed by HPLC. Experiments were conducted in triplicate; error bars show the standard deviation.

#### *General method for HiC-catalyzed depolymerisation of PLA under RAging conditions using 2 M glycine buffer*

The PLA powder (150 mg, 2.08 mmol) was placed in a 15 mL unsleeved PTFE milling jar containing one $ZrO_2$ ball (10 mm diameter). After the addition of commercial HiC enzyme (150 μL, 0.975 mg protein) and 2 M glycine buffer at pH 10 (525 μL, $\eta$ = 4.5 μl mg$^{-1}$), the mixture was milled hourly at 30 Hz for 5 minutes between periods of static incubation at 55°C. Samples (20-50 mg) were collected regularly for HPLC analysis. The experiments were performed in triplicate. The results are presented on **Figure S5**. The results of a similar mechanoenzymatic depolymerisation process mechanically activated just once are included for comparison.

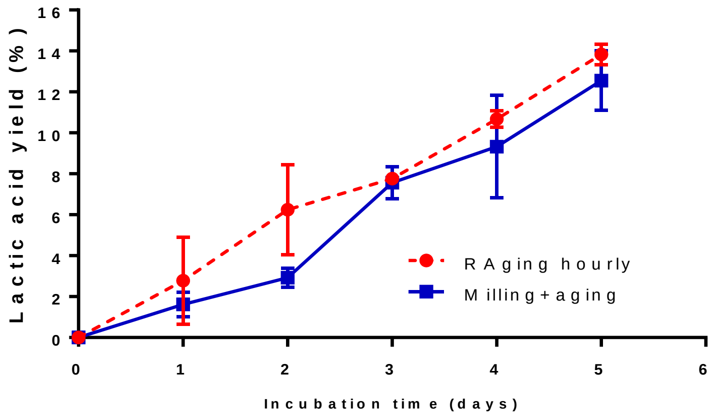


**Figure S5**. Effect of hourly milling (30 Hz, 5 minutes) on the yield of lactic acid for the HiC-catalyzed depolymerisation of PLA. The reactions were performed at $\eta$ = 4.5 μL·mg$^{-1}$ using 2 M glycine buffer at pH 10, with 150 mg of PLA, 0.65%w/w HiC, and static incubation at 55°. Experiments were conducted in triplicate; error bars show the standard deviation.

***PLA aminolysis catalyzed by HiC in the presence of high molarity buffer under moist-solid conditions***

A minor component of PLA aminolysis was observed during PLA depolymerisation under the following conditions. PLA powder (150 mg, 2.08 mmol) was placed in a 15 mL sleeved PTFE milling jar containing one $ZrO_2$ ball (10 mm diameter). After addition of the commercial enzyme preparation (150 μL, 0.97 mg protein) and glycine or Tris buffer at the desired pH ($\eta$ = 4.5 μl $mg^{-1}$, 525 μL), the reaction mixture was treated to ball milling at 30 Hz for 15 minutes, before static incubation at 55°C. Samples of the reaction mixture (15-40 mg) were collected for HPLC analysis. The experiments were performed in triplicate. Aminolysis products (**2** and **4** in main manuscript) were detected by HPLC.

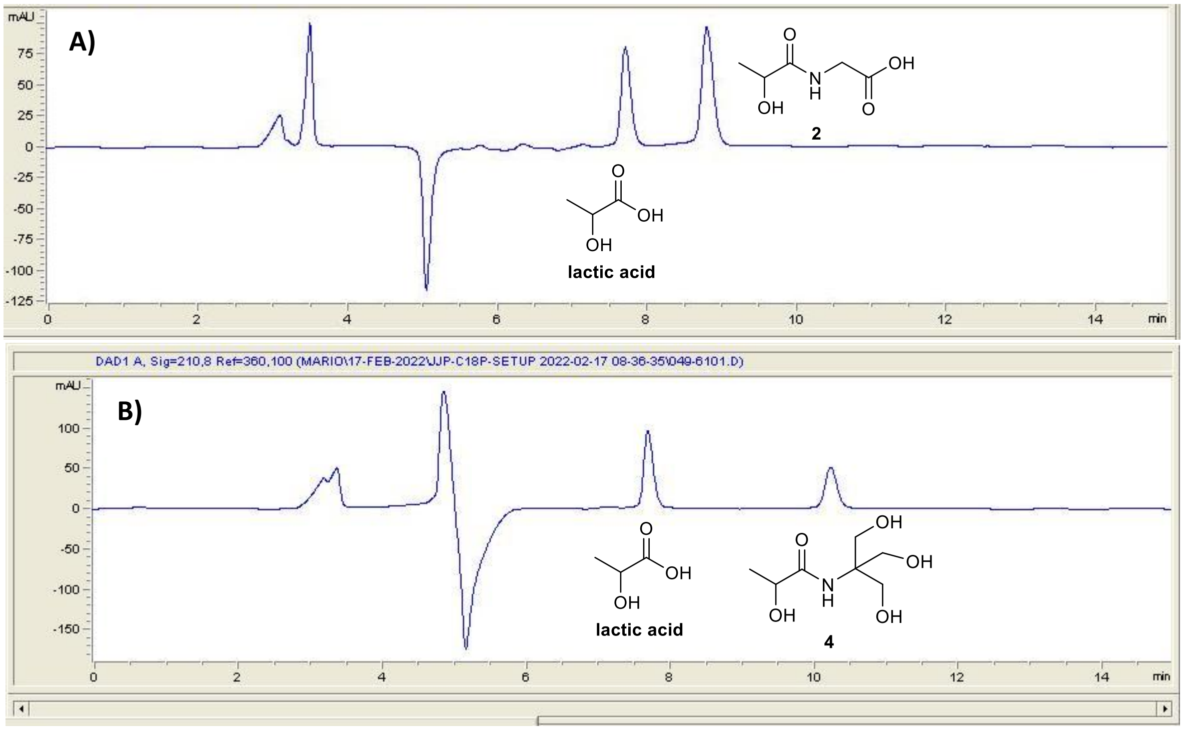


**Figure S6**. HPLC chromatograms of post incubated (5 days at 55°C) samples from the mechanoenzymatic depolymerisation of PLA using HiC (0.65% w/w) and A) 2 M glycine buffer pH 9, or B) 3 M Tris buffer pH 10 ($\eta$ = 4.5 μl $mg^{-1}$,see *Equipment* section for the detailed method). Lactic acid retention time: 7.4 min, amide product **2** retention time: 8.7 min, and amide product **4** retention time: 10.3 min.

Amide products **2** and **4** were isolated by semi-preparative HPLC, and recovered at a purity of 94% and 88%, respectively. These samples were used for further characterization and as standards for HPLC quantification (**Figure S7**).

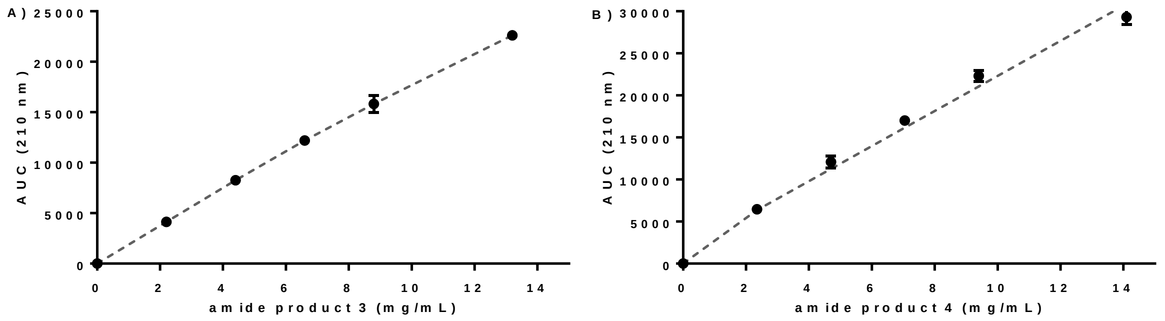


**Figure S7**. Calibration curve for amide products **2** and **4**, based on HPLC peak area (AUC). A) The goodness of fit for amide product **2** was $R^2$ = 0.9977, given the equation AUC = 1716.6 [**2**] + 434.74. B) The goodness of fit for amide product **4** was $R^2$ = 0.9868, given the equation AUC = 2083.8 [**4**] + 1472.

### ***Control experiments to rule out lactic acid as a promoter of PLA aminolysis under moist-solid conditions***

Control experiments were conducted under the following conditions. PLA powder (150 mg, 2.08 mmol) was placed in a 15 mL sleeved PTFE milling jar containing one $ZrO_2$ ball (10 mm diameter). After addition of the commercial enzyme preparation (150 μL, 0.97 mg protein), and either 2 M glycine buffer pH 10 containing 8.6 μL of fresh lactic acid or 3 M Tris buffer pH 9 containing 11.3 μL of fresh lactic acid ($\eta$ = 4.5 μl $mg^{-1}$, 525 μL), the reaction mixture was treated to ball milling at 30 Hz for 15 minutes, before static incubation at 55°C. The experiments were performed in triplicate.

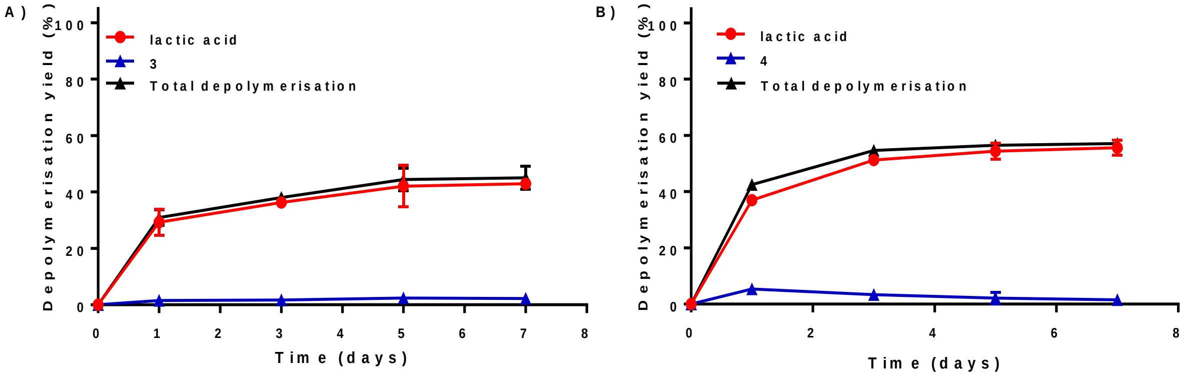


**Figure S8**. Effect of lactic acid on the aminolysis of PLA catalysed by HiC (0.65% w/w) under moist-solid reaction conditions ($\eta$ = 4.5 μL·$mg^{-1}$) measured in the presence of A) 2 M glycine buffer pH 10 containing 8.6 μL of lactic acid, or B) 3 M Tris buffer pH 9 containing 11.3 μL of lactic acid, after milling for 15 minutes at 30 Hz and static incubation at 55°C for variable periods. Experiments were conducted in triplicate; error bars show the standard deviation.

#### ***Spectrophotometric assay to quantify the protease activity of HiC using N-(4-nitrophenyl) butyramide as substrate***

The protease activity of HiC was quantified by spectrophotometric analysis of the 4-nitroaniline product (**Scheme S1**) at 405 nm ($\varepsilon$ = 9.96 $mM^{-1}cm^{-1}$). In the assay, 185 µL of a *N*-(4-nitrophenyl) butyramide (4-NAHx) solution (250 µM, 6% DMSO) and either 2 M glycine or 3 M Tris buffer at pH 9 or 10 was placed in a well of a 96-well plate and incubated at 45°C for 5 minutes before addition of 15 µL of HiC (Novozyme 51032; pre-incubated at 45°C). The reaction was monitored for 30 minutes (**Figure S9**) and the experiments were conducted in triplicate.

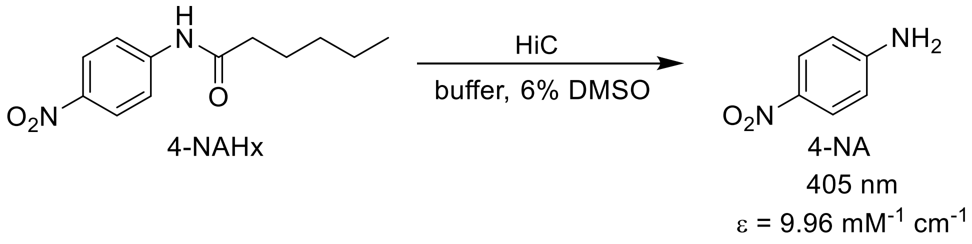


**Scheme S1**. Hydrolysis of *N*-(4-nitrophenyl) butyramide (4-NAHX) by HiC to produce 4-nitroaniline (4-NA) quantifiable at 405 nm.

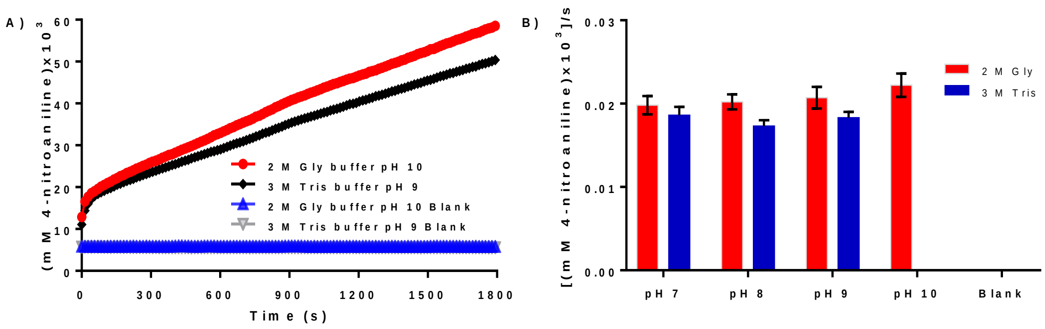


**Figure S9**. Hydrolysis of *N*-(4-nitrophenyl) butyramide by HiC (Novozyme 51032) using either 2 M glycine or 3 M Tris buffer at pH 9 or 10. Control experiments (blanks) were conducted using water instead of HiC. For clarity, the standard deviation was omitted in A), and the initial rate of a series of reactions are plotted in B).

#### ***General method for HiC-catalyzed depolymerisation of PLA with a second addition of buffer after 2 days***

The PLA powder (150 mg, 2.08 mmol) was placed in a 15 mL sleeved PTFE milling jar containing one $ZrO_2$ ball (10 mm diameter). After the addition of commercial HiC enzyme (150 µL, 0.975 mg protein) and either 2 M glycine buffer at pH 10 or 3 M Tris buffer at pH 9 (525 µL, $\eta$ = 4.5 µl $mg^{-1}$), the mixture was milled at 30 Hz for 15 minutes, before static incubation at 55°C for 2 days. More buffer (225 µL, for a total of $\eta$ = 6 µL/mg) was then added. Samples of the reaction mixture (15-40 mg) were collected for HPLC analysis. Experiments were performed in triplicate. Control experiments without

HiC were included. Control experiments adding MilliQ® water instead of buffer were also performed (**Figure S10**).

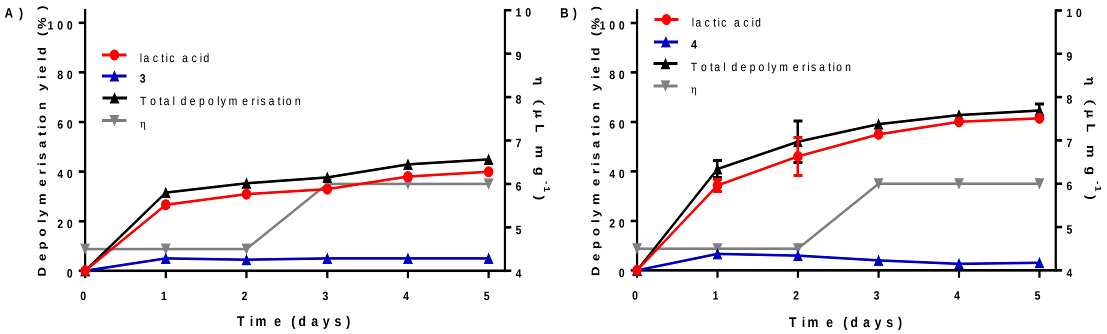


**Figure S10**. Yield of total depolymerisation, or of lactic acid, obtained when the addition of a second batch (2 days of incubation) of buffer was replaced by MilliQ® water, for the HiC-catalysed (0.65% w/w) hydrolysis of PLA treated to milling for 15 minutes at 30 Hz and static incubation at 55°C, using A) 2 M glycine buffer pH 10, or B) 3 M Tris buffer pH 9 and the initial buffer. Experiments were conducted in triplicate; error bars show the standard deviation.

#### *Chemoenzymatic synthesis of rac-5 from PLA*

Compound *rac*-**5** was synthesized from PLA in a two-step chemoenzymatic process (**Scheme 1**, main manuscript). The first step was an HiC-catalyzed depolymerisation of PLA to lactic acid in a moist-solid mixture. Thus, the PLA powder (300 mg, 4.16 mmol) was placed in a 15 mL sleeved PTFE milling jar containing one $ZrO_2$ ball (10 mm diameter). After addition of the commercial enzyme preparation (300 μL, 1.95 mg protein) and 3 M Tris buffer at pH 9 (1050 μL, $\eta$ = 4.5 μl $mg^{-1}$), the reaction mixture was milled at 30 Hz for 15 minutes. The mixture was next transferred into a 5 mL Eppendorf tube, sealed with parafilm, and incubated at 55°C for 2 days. After which 450 μL of fresh 3 M Tris buffer at pH 9 was added before further incubation at 55°C for 3 days. The product was collected by washing the mixture with water twice using centrifugation (6700 x g) to separate the layers. The lactic acid content was quantified by HPLC, yielding quantitative depolymerisation with minimal aminolysis (<0.1%).

In the second step, this crude lactic acid/lactate was reacted with *o*-phenylenediamine to produce *rac*-**5**. This was achieved with the addition of HCl (3 ml of 6 N) to the crude lactic acid (3.35 mmol), stirring for 15 minutes, addition of *o*-phenylenediamine (361 mg, 3.35 mmol) and heating under reflux for 6 hours. The product *rac*-**5** was a brown/red solid obtained in 60% overall yield (NMR yield), based on the initial amount of PLA.

***Characterization of lactic acid, amides products rac-2 and rac-4, and rac-5***

***Lactic acid:*** $^{1}H$-NMR (300 MHz, $D_2O$) δ: 3.91 (q, 1H, *J* = 6 Hz), 1.13 (d, 3H, *J* = 6 Hz). $^{13}C$-NMR (75 MHz, $D_2O$) δ: 182.50, 68.42, 20.11. Values are consistent with those of commercial lactic acid.

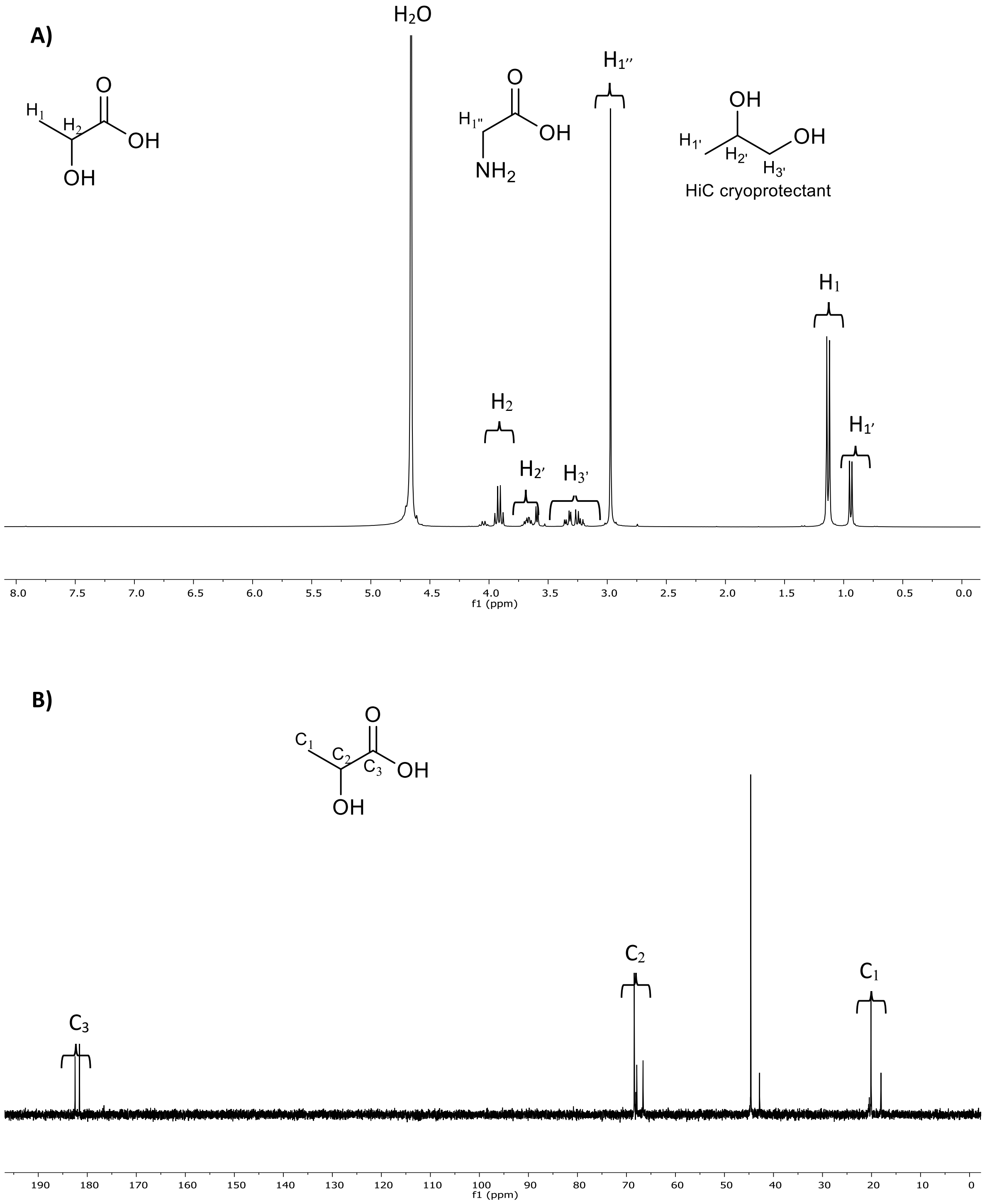


**Figure S11.** $^{1}H$ A) and $^{13}C$ B) NMR spectra (in $D_2O$) of the reaction mixture after mechanoenzymatic PLA depolymerisation under optimal conditions using 2 M glycine buffer pH 10.

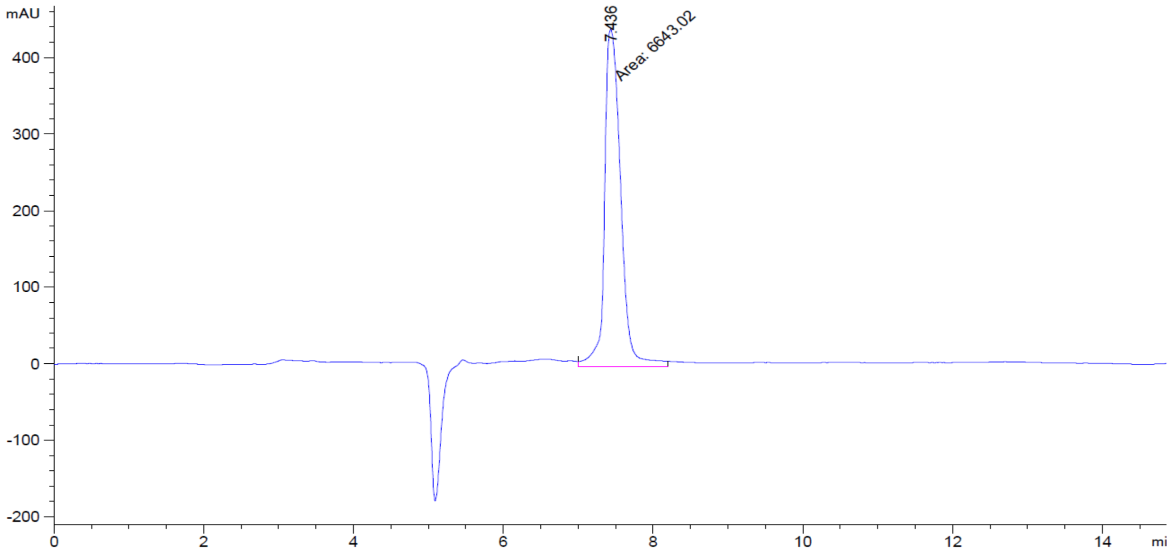


**Figure S12**. HPLC chromatogram of commercial lactic acid (10 mg/mL, 90% purity, see *Equipment* section for the detailed methods).

***rac*-2**: $^{1}H$-NMR (300 MHz, $D_2O$) δ: 4.27 (q, 1H, *J* = 6 Hz), 3.9 (s, 2H), 1.35 (d, 3H, *J* = 6 Hz). $^{13}C$-NMR (75 MHz, D2O) δ: 177.05, 174.47, 67.59, 41.45, 19.35. HRMS (ESI$^{+}$) *m/z* $[C_5H_8NO_4Na_2]^+$ calcd: 192.0243, found: 192.0241.

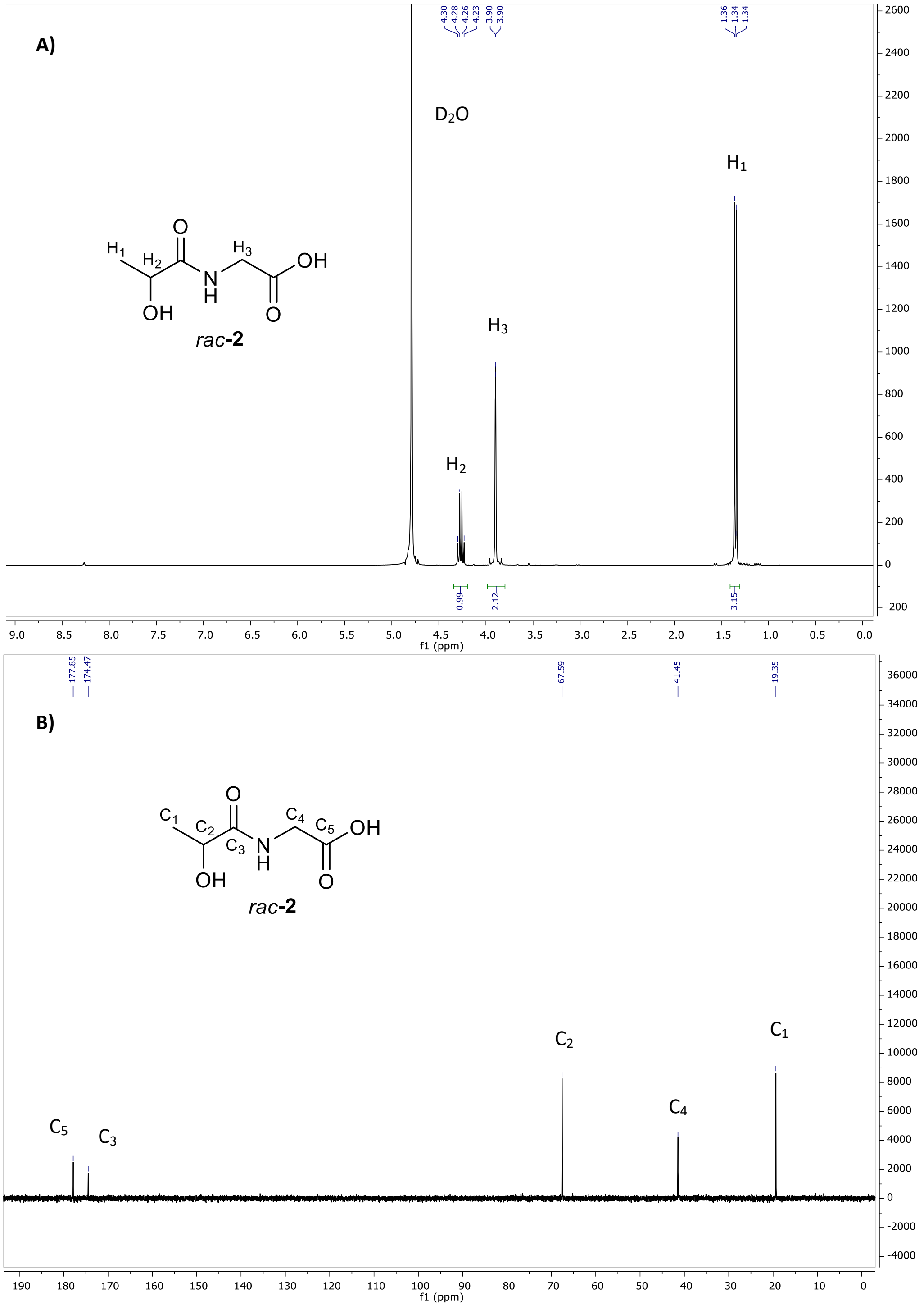


**Figure S13**. $^{1}H$ A) and $^{13}C$ B) NMR spectra of ***rac*-2** in $D_2O$.

***rac*-4**: $^{1}$H-NMR (300 MHz, $D_2O$) $\delta$: 4.05 (q, 1H, *J* = 6 Hz), 3.59 (s, 6H), 1.18 (d, 3H, *J* = 6 Hz). $^{13}$C-NMR (75 MHz, D2O) $\delta$: 177.54, 67.92, 61.22, 60.38, 19.52. HRMS (ESI$^{+}$) *m/z* $[C_7H_{15}NO_5Na]^{+}$ calcd: 216.0842, found: 216.0844.

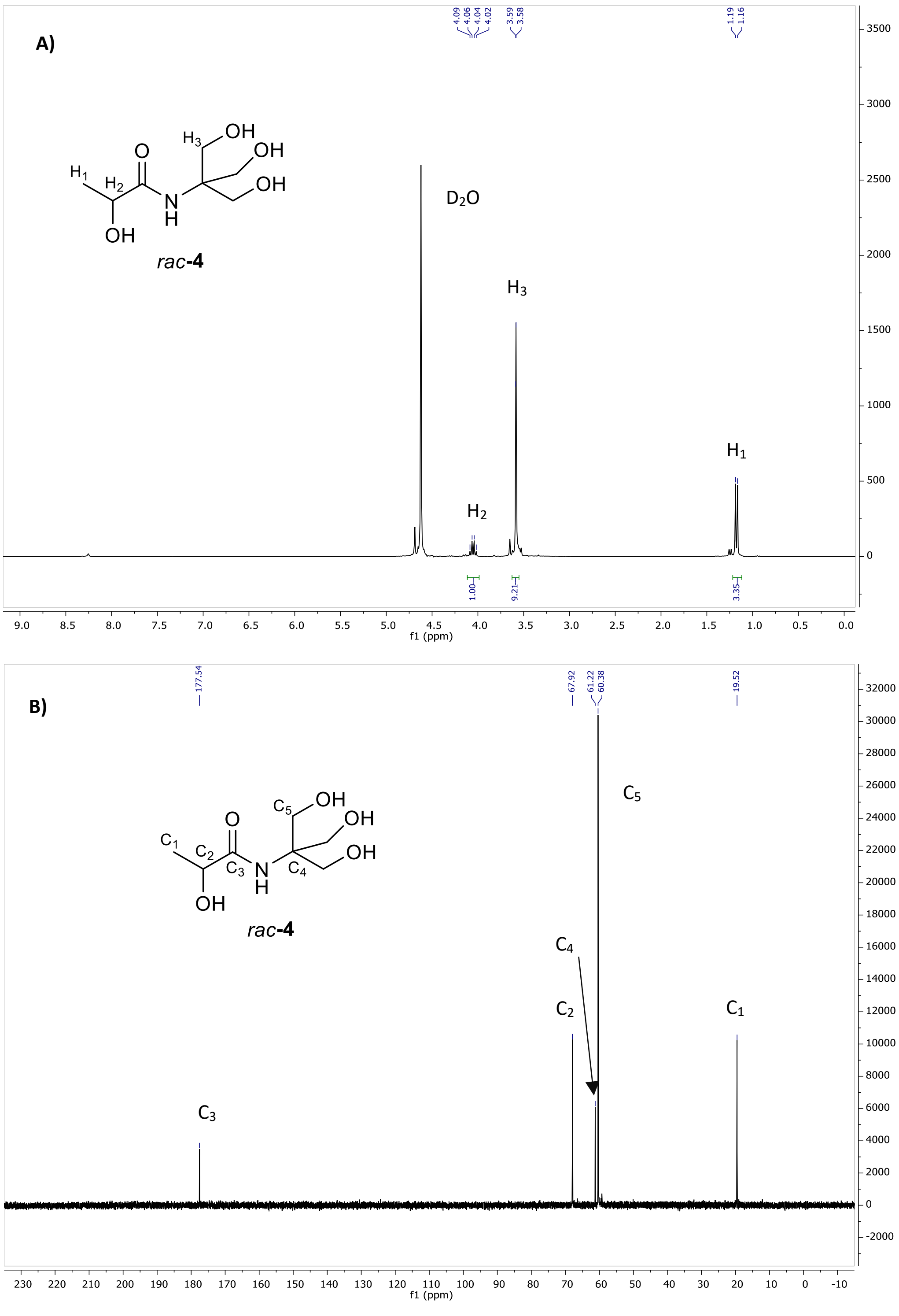


**Figure S14**. $^{1}$H A) and $^{13}$C B) NMR spectra of ***rac*-4** in $D_2O$.

***rac*-5**: $^{1}$H-NMR (300 MHz, MeOH-*d*4) δ: 7.53 (m, 2H), 7.20 (m, 2H), 5.07 (q, 1H, *J* = 6 Hz), 1.62 (d, 3H, *J* = 6 Hz). $^{13}$C-NMR (75 MHz, MeOH-*d4*) δ: 158.4, 137.9, 121.9, 114.4, 64.1, 21.8. HRMS (ESI$^{+}$) *m/z* $[C_9H_{11}N_2O]^+$ calcd: 163.0866, found: 163.0870.

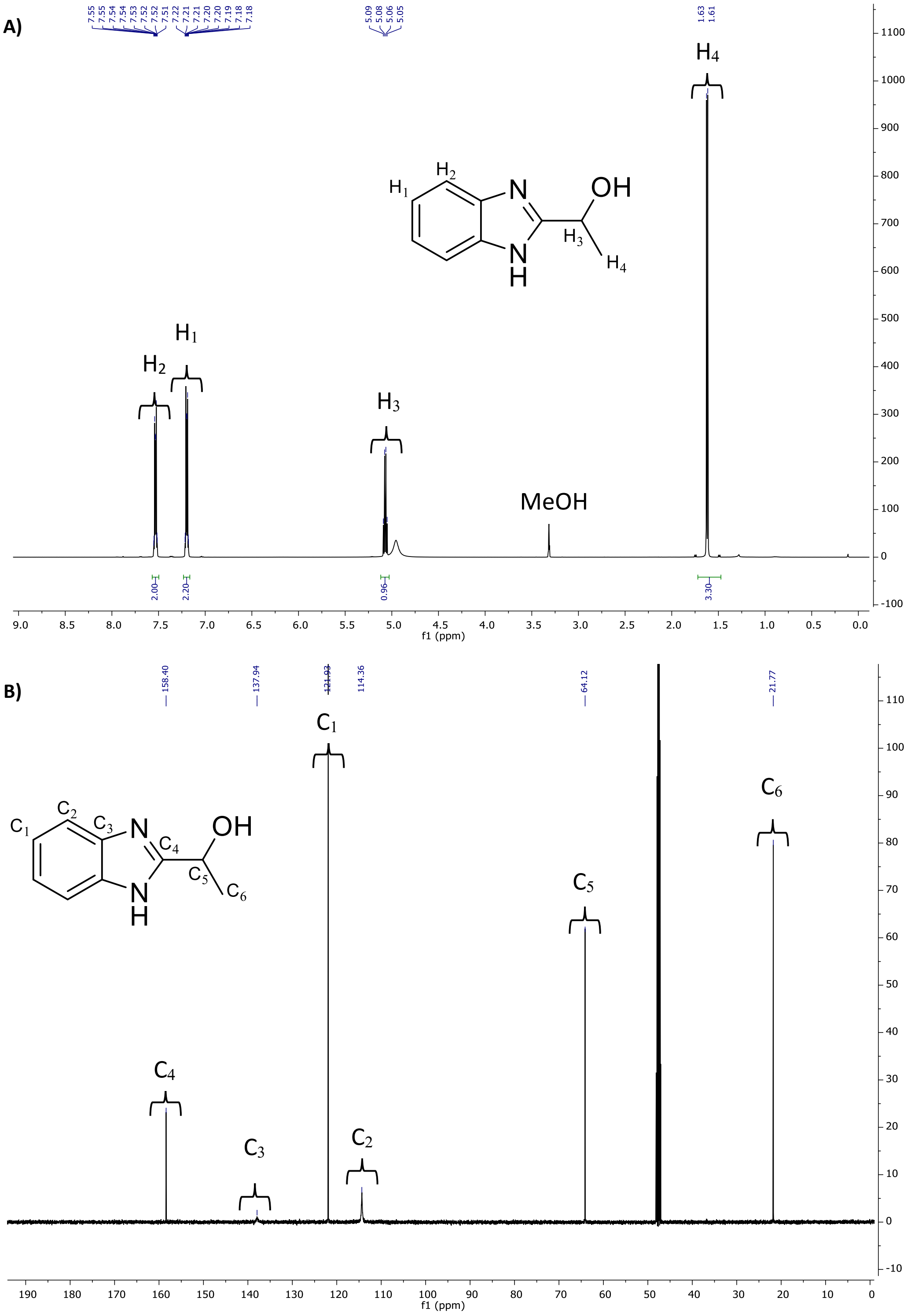


**Figure S15**. $^{1}$H A) and $^{13}$C B) NMR spectra of ***rac*-5** in MeOH-*d*4.

**Table S1**. Reported PLA degradation experiments mimicking natural biodegradation conditions (adapted from *Ref*. 1)

| Environment | Controlled conditions | Scale | Measurement method | Biodegradation (%) | Period of biodegradation (Days) |
|---|---|---|---|---|---|
| Soil | 30% moisture | Buried in alluvial-type soil (12-15 cm) | Weight loss | 10 | 98 |
| | 25°C, 60% humidity[a] | Mixed in 50 g soil | Weight loss | 13.8 | 28 |
| | 30°C, 80% humidity[b] | Buried in topsoil-outside | Weight loss | 37.4 | 56 |
| | 30°C, 80% humidity[c] | Buried in topsoil-lab | Weight loss | 43 | 56 |
| | 80% moisture[d] | Buried in alluvial-type soil (12-15 cm) | Weight loss | >60 | 98 |
| Compost | 58°C | Lab-scale compost reactor (1L bottle) | Produced $CO_2$ | 13 | 60 |
| | 65°C, pH 8.5, 63% humidity | Wooden box | Produced $CO_2$ | 84 | 58 |
| | 55°C, 70% moisture | Lab-scale composting setup | Produced $CO_2$ | 70 | 28 |
| | 58°C, 60% humidity, aerobic | Lab-scale | Weight loss | 60 | 30 |
| | 58°C, aerobic[e] | Polypropylene reactor | Weight loss | 63.6 | 90 |
| | 58°C[e] | Laboratory scale plastic reactor | Weight loss | 100 | 28 |
| Aquatic environments | 25°C, 16h light and 8h dark[f] | Lab-scale | Weight loss | <2 | 365 |
| | 25°C, 16h light and 8h dark[g] | Lab-scale | Weight loss | <2 | 365 |
| | 30°C[h] | Lab-scale | Produced $CO_2$ | 3.1-5.7 | 180-365 |

[a]Powdered PLA was used. [b]Mixture of PLA/NPK fertilizer (63.5/37.5). [c]Mixture of PLA/NPK fertilizer/EFB compost (25/37.5/37.5). [d]Mixture of PLA/sisal fiber (60/40). [e]Synthetic material containing compost. [f]Freshwater. [g]Sea water.[h]Marine environment.

**Table S2**. Reported biocatalytic (with microorganisms or enzymes) PLA depolymerisation methods with quantification of the yield of PLA biodegradation or depolymerisation, starting from a similar PLA substrate as the one used in this manuscript.*

| Biocatalyst | PLA source | Conditions | Measurement method | Depolymerisation (%) | Duration (Days) |
|---|---|---|---|---|---|
| *Actinomadura* sp.[2] | Emulsified PLA | 100 mM Tris-HCl buffer pH 9, 60°C | Turbidity | - | - |
| *Bordetella petrii*[3] | PLA powder | Compost, 37°C | $CO_2$ | 68 | 40 |
| Bacillus licheniformis[4] | PLA composite | 32°C | Weight loss | 90 | 150 |
| *Pseudonocardia* sp. RM423[5] | Emulsified PLA | 30°C | Weight loss | 15 | 28 |
| *Trichoderma viride*[6] | PLA composite | 28°C | Weight loss | 12 | 21 |
| Cutinase enzyme from *Aspergillus oryzae*[7] | Emulsified PLA | 10 mM Tris-HCl buffer pH 8, 37°C | Turbidity | Active | - |
| Cutinase enzyme from *Cryptococcus* sp.[8] | Emulsified PLA | 20 mM Tris-HCl pH 7, 30°C | Turbidity | >90 | 2.5 |
| Depolymerase from *Pseudomonas tamsuii* TKU015[9] | Emulsified PLA | 50 mM phosphate buffer pH 7, 50°C | Lactic acid kit | Active | - |
| PLA-degrading enzyme from *Laceyella sacchari* LP175[10] | Emulsified PLA | 100 mM Tris-HCl buffer pH 9, 60°C | Turbidity | Active | - |
| PLA-degrading enzyme from *Amycoltopsis orientalis*[11] | PLA | 0.1 M sodium phosphate buffer pH 7, 0.5% (w/v) octyl glucopyranoside, 30°C | TOC | Soluble products | 8 hours |
| Proteinase K[12] | Emulsified PLA | 50 mM buffer pH 8.6, 40°C | Turbidity | Active | - |
| Esterase from *Pseudomonas aeruginosa*[13] | PLA film | Minimum medium, 30°C | SEM | Active | 28 |
| Lipase enzyme from *Sphingobacterium* sp.[14] | PLA film | 100 mM Tris HCl pH 8, 37°C, 500 rpm | LCMS | 27 | 3 |
| ABO2449 from Alcanivoraz borkumensis[15] | Emulsified PLA | 50 mM Tris-HCl buffer pH 8, 30°C | Turbidity/HPLC | >90 | 2 days |

*For extensive reviews see *Ref*. 16-18.
